# Structure of ABCB1/P-glycoprotein bound to the CFTR potentiator ivacaftor

**DOI:** 10.1101/2021.06.11.448073

**Authors:** Alessandro Barbieri, Nopnithi Thonghin, Talha Shafi, Stephen M. Prince, Richard F. Collins, Robert C. Ford

## Abstract

ABCB1 (P-glycoprotein) is an ATP binding cassette transporter that is involved in the clearance of xenobiotics and it affects the disposition of many drugs in the body. Here we have studied ABCB1 in the drug-bound and drug-free states, simultaneously, using high contrast cryo-electron microscopy imaging and a Volta phase plate. The binding of the potent CFTR potentiator, ivacaftor, at a site in the central aqueous cavity is mediated by transmembrane α-helices 3,6,10,11 & 12. Binding is associated with a wider separation of the two halves of the transporter in the inward-facing state. Induced-fit changes the nucleotide binding domains in a way that may explain their increased affinity for ATP when drug is bound. Comparison of ivacaftor-bound structures of CFTR and ABCB1 suggests common features in the binding modes.

## Introduction

ABCB1 is an ATP-dependent multi-drug transporter involved in active efflux of many drugs and xenobiotics from cells(1). Overexpression or activation of ABCB1 can lead to multidrug resistance in cancer cells(2–5). Drugs are thought to bind to the inward-facing conformation of ABCB1(6,7), which is more favoured in the post hydrolytic or nucleotide free conditions(8–10). The drug is then translocated across the plasma membrane by transition of the protein to the outward facing state, which is more favoured when ATP is bound(10). The dynamic conformational nature of ABCB1 (and other ABC family members) has challenged structural studies by X-ray crystallography and cryo-electron microscopy (cryo-EM). Stabilising antibodies and ATP hydrolysis-inactivating mutations have been used previously to generate ∼4 Å resolution structural data by cryo-EM(7,11,12). Similarly, trapping the protein in a post-hydrolytic state with vanadate and ATP yielded an ∼8Å resolution structure (13). However these approaches could be criticised because the protein was in an artificial, stabilised state and under these experimental conditions, allosteric changes may be altered or inhibited. In this report we describe studies of ABCB1 at the early stages of its transport cycle (both apo- and drug-bound states) in the absence of any stabilising antibodies and without inactivation or trapping.

Recent studies have demonstrated that the novel cystic fibrosis (CF) therapeutic, ivacaftor, is a transported substrate of ABCB1(14,15). This drug is a potentiator of the channel activity of an ABCB1 homolog, CFTR/ABCC7 (16), and ivacaftor is now widely used in the clinic, both on its own as well as in combination with CFTR-corrector compounds such that roughly 90% of CF patients may now be treatable(17,18). A prior structure for ivacaftor bound to CFTR showed the drug on the external surface of the membrane-spanning region and within the hydrophobic region of the detergent micelle(19). The drug itself is highly hydrophobic and was used at a relatively high concentration, but mutagenesis studies and experiments with another potentiator drug with slightly different structure were undertaken to provide further evidence that the position of the drug in CFTR represented a common binding mode. A further interesting observation in the study of ivacaftor binding to CFTR was that there was minimal conformational change upon drug binding. Hence there was no induced fit involved in the protein, and because of the rigid nature of the drug, conformational selection (of the drug) may not occur. Structure activity relationships (SAR) of ivacaftor homologues with potentiating and inhibiting properties identified the quinoline nitrogen as crucial for activity via hydrogen bonding. π-π stacking with nearby residues by the same quinoline group was also found to be important. The planar nature of the drug was also shown to be a significant factor in the SAR study. The hydroxyl group of the hydroxyphenyl moiety is polar with a computed pKa of 11 (CHEMBL2010601) and points towards the guanidino group of R933 in the structural model (16,19), implying polar interactions that are too distant (6 Å) for hydrogen bonding.

In this study we have examined the binding of ivacaftor to ABCB1. In order to minimise non-specific binding of the hydrophobic drug, we employed a concentration likely to be encountered *in vivo* (20), and at a sub-stoichiometric ratio with the protein. This approach had the advantage of allowing both drug-bound and drug-free forms of the protein to coexist as a mixture in the protein/drug solution immediately prior to flash freezing and cryo-electron microscopy. We reasoned that if drug binding caused an induced fit and a significant conformational change in ABCB1, we should be able to distinguish different conformations using 3D classification of the particles. If no induced fit occurred (as was the case for CFTR(19,21)), we expected to obtain a single three-dimensional structure but with fractional occupancy of the drug at its binding site, leading to weak density for ivacaftor in the experimental 3D density map. A third possibility existed: that drug binding would cause conformational changes, but that discrimination of these different conformations would not be possible by the image processing routines. In this case we would expect a smeared, low resolution three-dimensional map for the protein with no information about drug binding. In order to minimise the chances of the latter scenario we employed the Volta phase plate to collect very high signal:noise data with minimal under focus. There are difficulties in the use of the Volta phase plate that restrict its current wide application (21) and there are currently only a few examples of studies resulting in resolution better than 3Å (21). Nevertheless we decided that the high contrast provided by the Volta phase plate would allow us to distinguish different conformational states of ABCB1, and that even at 4 to 5 Å resolution, sufficient information on drug binding could be obtained to determine the ivacaftor binding mode in ABCB1.

## Results

### Ivacaftor binding affinity of murine ABCB1

Ivacaftor has been identified as a competitive inhibitor for human ABCB1 (14,15). Ivacaftor stimulated human ABCB1 ATPase activity eight-fold, suggesting it could also be considered as a transported allocrite and its dissociation constant (Kd) was estimated at 0.3 μM using a fluorescent dye transport assay with the purified protein (14). Similarly, a Kd of 0.2 μM was estimated using a cell-based assay (15). We tested ivacaftor binding to murine ABCB1 using various methods: Static light scattering and tryptophan fluorescence changes yielded Kd values of 0.2 μM (static light scattering) and ≅1 μM (tryptophan fluorescence) (Supplementary Figure 1a,b). The binding of a sulfhydryl-reactive fluorescent dye (7-Diethylamino-3-(4′-Maleimidylphenyl)-4-Methylcoumarin, CPM) to solvent-exposed ABCB1 cysteine residues detected an unfolding transition between 30-50°C that was sensitive to ivacaftor with a Kd estimated at 0.2 μM (Supplementary Figure 1c). Hence we concluded that the affinity for ivacaftor of murine ABCB1 was similar to human ABCB1. Moreover the detection of light scattering changes upon ivacaftor binding implied conformational changes in ABCB1 (e.g due to an induced fit). Cryo-EM studies were subsequently performed with a concentration of ivacaftor (2 μM) that was well above the Kd, although at this level it was still at a sub-stoichiometric ratio versus protein (estimated at 1.1 mg/ml, or roughly 8 μM using a molecular mass of 150kDa for ABCB1).

### Cryo-EM of ivacaftor-ABCB1 complexes

Images (2436) were recorded with the Volta phase plate, and after initial rejection of poor quality images by eye, the contrast transfer function (CTF) of each was estimated. A total of 1537 images gave CTF fits that were judged to be reliable whilst 868 were rejected at this stage. A second criterion was applied to eliminate all images affected by excessive defocus values, and a total of 131 further images were eliminated, with a final set of 1406 images (Supplementary Figure 2). As expected, the ABCB1 particles were readily identifiable in the images in the inward-facing conformation (Supplementary Figure 3) and automated particle picking was performed. Reference-free classification of the projections of the resulting picked particles allowed a further pruning of the dataset by deleting any non-ABCB1 contaminants as well as small ABCB1 aggregates (Supplementary Figure 4). Several iterations of 2D classification were completed to remove bad particles and this was judged to be complete when consistent 2D classes were obtained. The reproducibility of the 2D classification was then assessed by splitting the dataset into two roughly equally populated sub-sets which individually showed the same outcome (Supplementary Figure 2).

A*b-initio* 3D classes of the final ABCB1 dataset were generated from a subset of ∼50,000 particles and these are shown in Supplementary Figure 5. At this early stage, the 3D classes were relatively low resolution (∼7 to 8 Å, as estimated by Resmap(22)), nevertheless, the two procedures clearly identified only inward-facing conformations, even when five classes were allowed. Sets of particles displaying a wider separation of the NBDs (classes *a* & *c*) were also picked out by the image processing package. For each of the five *ab-initio* maps a PDB model was derived using Molecular Dynamics Flexible Fitting (*see Methods*) and results were compared. Utilising the center of mass of the NBDs as a reference, classes *b*, *d* and *e* showed a narrower separation of the NBDs (from 56Å to 53 Å) while both classes *a* and *c* were found to have a wider distance (60 Å). A moderate low-pass filter was applied to the *ab-initio* maps to avoid over-fitting effects. CTF parameters were also further refined at this stage with per-micrograph CTF fits using the dedicated *cis*TEM functionality for this. Refinement of the full dataset (104,000 particles) starting with each of the five *cis*TEM *ab-initio* 3D classes was carried out with a combination of auto- and manual-refinement, and an atomic model was generated for each resulting refined 3D map using the PHENIX *real space refinement* routine (23). The 3D maps, compared at a density threshold yielding a map volume of 153,000 Å^3^ are displayed in Figure 1. The resolution of each was assessed by splitting each data set in half and separately refining each half dataset. The correspondence between half-maps was analysed in reciprocal space using Fourier shell correlation (FSC) as well as using the Resmap routine, which calculates local resolution in the maps (22,24). The resolution assessments as well as other data pertaining to the five refined maps and the five atomic models are summarised in Table I. Each 3D map could be generated from fewer particles than in prior studies of ABCB1 by cryoEM (7,12,13), due to the high contrast of individual particle projections produced with the Volta phase plate. The resolution achieved was estimated at 4-6Å for all the maps (Table I, Supplementary Figures 6-8). Map resolution was not correlated with particle numbers, consistent with the idea that, for structural studies of ABCB1, intrinsic protein flexibility is likely to be a major factor in determining the resolution achievable. Interestingly, map *a* displayed the lowest resolution of all the maps in the various tests employed, although the overall map-to-model correlation was better (Table 1, Supplementary Figure 8). The Resmap analysis of map *a* implied that variation between the half-maps was mainly confined to the periphery of the NBDs and the micelle (Supplementary Figure 6). Conversely, map *e*, with the closest approach of the NBDs, was estimated to have the highest resolution by the Resmap algorithm (22) and by Fourier shell correlation (24) (Supplementary Figure 7). In general, the map-to- model correlation was lower for the NBDs compared to the TMDs in all maps (Supplementary Figure 7).

**Figure 1:**
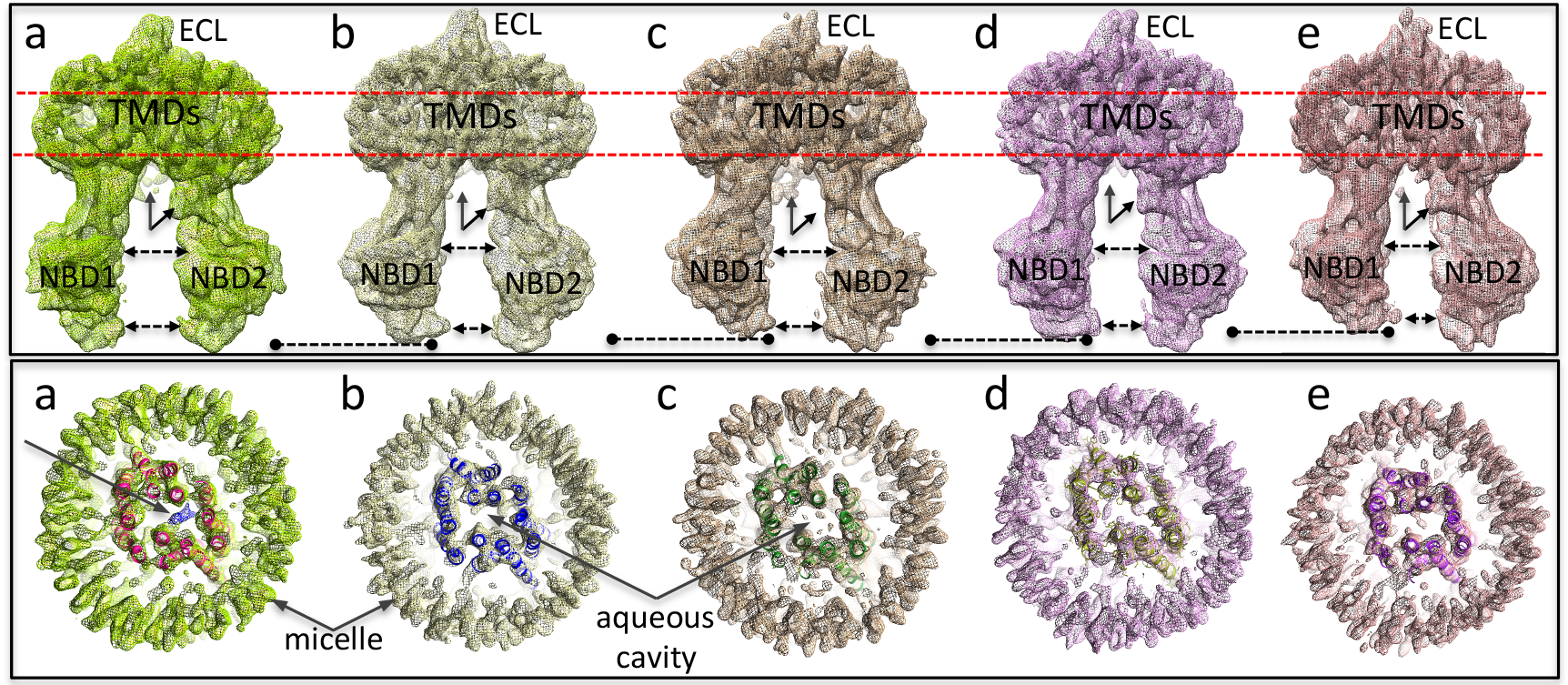
Differentiation of the five 3D maps (a-e). Maps are displayed at a density level enclosing a volume of 153,000 Å^3^. Upper panel: (1) NBD1 rotates and hinges upwards towards the TMDs; with this increasing from maps (a) through to (e), (dashed line with circles indicates the lower edge of NBD1). (2) NBD1-NBD2 separation, (dashed lines with arrowheads) is greatest for map (a) and (c), least for map (e) and intermediate for maps (b) and (d). (3) Density for the N-terminus (rearwards, vertical arrow) and the end of the NBD1-NBD2 linker (slanted arrow, front) is present in map (a), whilst other maps lack the N-terminus density (maps (b) and (d)) or lack the linker density (c). (4) The density for the extracellular loop (ECL) between TM helices 1 and 2 is weaker for maps (c) and (e). Lower panel: Slices through the transmembrane regions of the maps (as indicated by the red dashed lines in the upper panel). Real-space refined atomic models that were generated are also displayed using ribbon representation in contrasting colours. The diameter of the micelles in maps (a) and (c) are larger; map (e) has the smallest micelle with some distortion from circular. Map (a) contains a central density (blue mesh, arrow) not accounted for by the real-space refined model. Other maps have only small features in the aqueous central cavity. Map (e) shows the most compact transmembrane organization.

**Table I:**
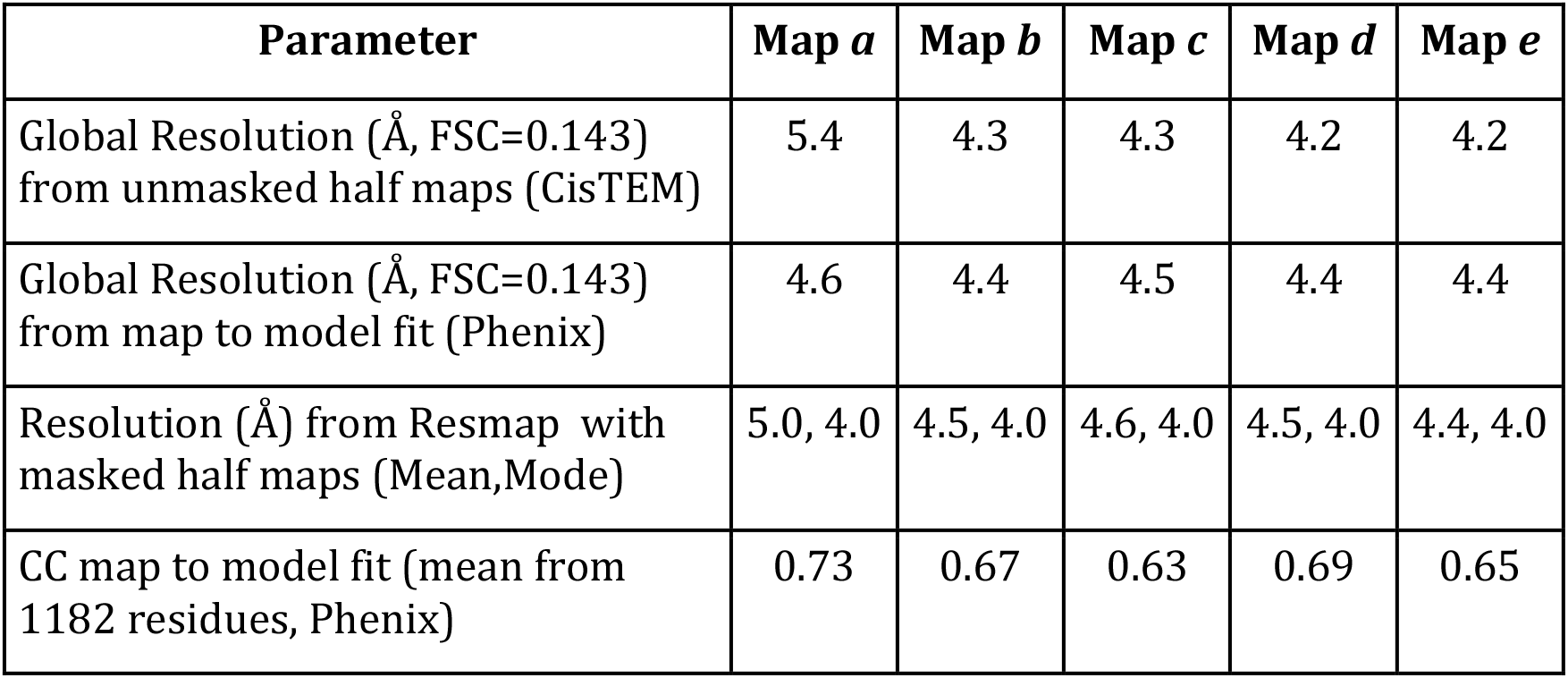
Summary of all the data for reciprocal space resolution assessment of the five maps as well as the real-space correlation coefficient (CC) between the atomic model and the relevant map.

### Interpretation of the 3D maps

The various 3D maps showed the typical hinging of the two halves of the molecule, giving differences in NBD separation, but it was clear that NBD1 flexing relative to the other ABCB1 domains was also being differentiated by the 3D classification algorithm: NBD1 appears to progressively rotate upwards towards the TMDs when one examines maps *a* through *e*; (Figure 1, upper panel). This movement can be compared to that of the shovel of a mechanical digger. Separation of NBD1 from NBD2 was greatest for maps *a* and *c*; was narrower in maps *b* and *d*; whilst map *e* had the closest approach of the NBDs (Fig. 1, upper panel). As previously reported (13), densities assigned to the N-terminus and to the end of the NBD1-NBD2 linker on the opposite side of the molecule can be observed in the maps, especially map *a (Fig. 1 upper panel)*. Maps *b* and *d* showed additional density for the linker, but lacked density for the N-terminal region. In contrast, map *c* showed the latter but appeared to lack the linker density. The density for the extracellular loop (ECL) between TM helices 1 and 2 was weaker for maps *c* and *e*, perhaps representing increased disorder in this loop which may be core glycosylated in the expression system used.

The transmembrane regions (Figure 1, lower panel) also showed some subtle differentiation between the five 3D maps. The diameter of the micelles in maps *a* and *c* are noticeably larger; whilst map *e* has the smallest micelle, displaying some distortion from a circular annulus. These differences may be a reflection of the degree of separation of the two halves of the protein, which is greatest for maps *a* and *c* and least for map *e*. The latter map shows the most compaction of the transmembrane α-helices.

### Drug binding site(s)

The transmembrane regions of all five maps (Figure 1, lower panel) displayed the large central aqueous cavity which connects to the cytoplasm as well as to the lipid bilayer (via the two lateral gaps formed between TM helices 4&6 and between 10&12). These lateral gaps are a common feature of type IV ABC transporters such as Sav1866, MsbA and ABCB1(25). In ABCB1 the central aqueous cavity has been found to house the binding sites for transported drugs and inhibitors (7,26,27). We therefore closely examined this cavity for evidence of ivacaftor binding in the five maps. The criteria we used were that: (i) any features due to ivacaftor should be comparable to protein in terms of their density level and (ii) that at this density level, the feature(s) should be roughly rod-shaped and (iii) have a continuous volume equivalent in size to that expected for the drug. Finally, (iv) once fitted, the ivacaftor molecule should not have serious clashes with nearby residues in the ABCB1 atomic model (i.e. none that could not be remedied with rotamer adjustment). Using these criteria, only map *a* contained a single feature consistent with ivacaftor (blue mesh, arrow, Figure 1, lower panel). All the other maps showed small features in the aqueous central cavity (Fig. 1 lower panel), but none of these met the first three criteria.

### Ivacaftor binding site

The additional density in the aqueous cavity of map *a* has a rocket-shape with two fin-like protrusions at one end. At the opposite end to the protrusions, the density is in close contact with TM6 and is close to F339 and Q343 (Figure 2). A single ivacaftor molecule could be fitted into this density, and the overall rocket shape allowed the discrimination of alternative 180°-rotated versions of the fit, with the 2,4 *tert-*butyl groups at one end of the ivacaftor molecule being fit to the two ‘fins’ of the rocket-shaped density. With this configuration, the drug is surrounded by residues contributed by TM helices 6, 10 & 12, and hence is positioned asymmetrically in one lobe of the aqueous cavity that is primarily formed by TM helices 1,2,3,6,10,11,12. Residues F339 and Q343 within TM6 appear to be close enough to form π-π (F339) and H-bonding (Q343) interactions with the quinoline group of ivacaftor (Figure 2, Figure 3a). Interestingly, the phenolic hydroxyl group also forms H-bonding with the close carboxylate of E871 within TM10, therefore contributing to the stabilization of the opposite end of ivacaftor. Small (glycine) residues at 868 and 985 in TM helices 10 and 12 may also be important: bulky amino acid side-chains at these positions would prevent accommodation of the *tert*-butyl groups of ivacaftor. I864, M872, L875 in TM10 and A981 in TM12 may also satisfy hydrophobicity requirements for the *tert*-butyl groups. (Figure 2, Figure 3a). Interestingly, K185 in TM3 extends up into the central cavity to within ∼10Å of the phenolic hydroxyl group of ivacaftor in a manner akin to R933 in human CFTR (19). Although the pKa of the phenolic hydroxyl group is predicted to be around 11, the stabilizing presence of the lysine residue in ABCB1 (or R933 in CFTR) may increase the probability of de-protonation. Hence partial charge-charge interactions may be formed by the side-chain amino group of K185 in the aqueous cavity.

**Figure 2:**
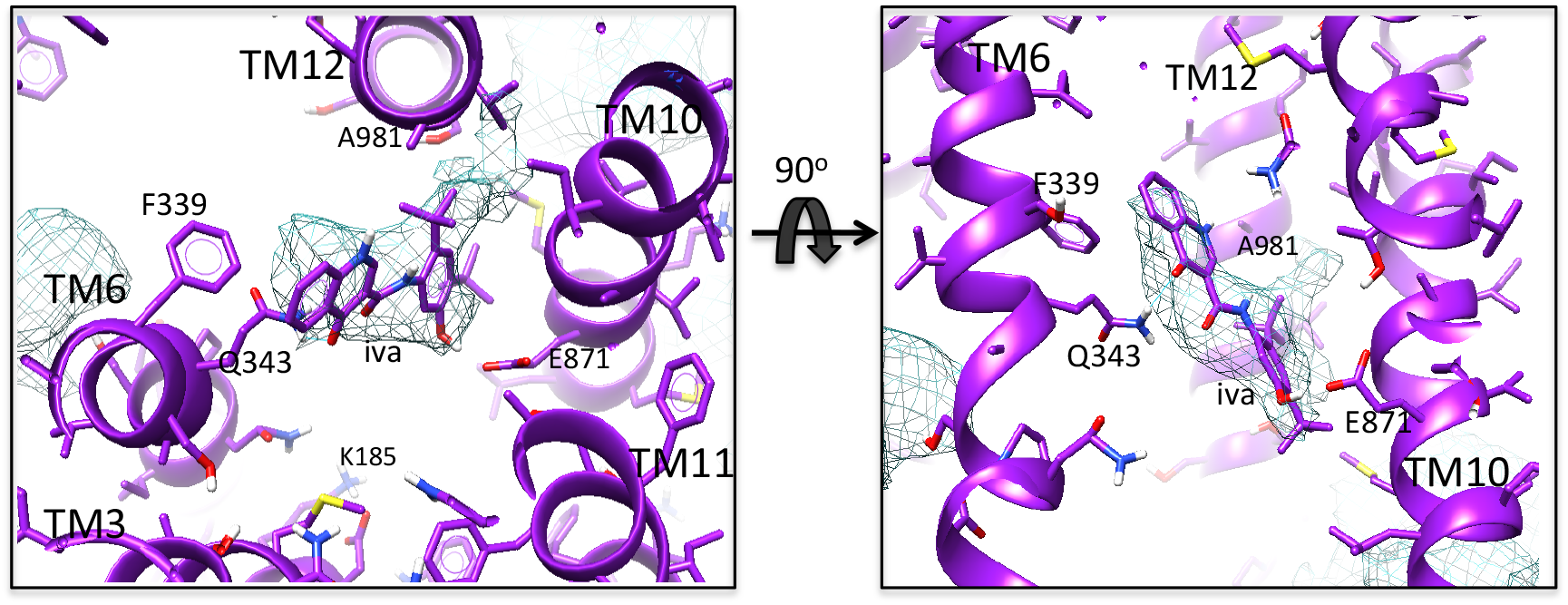
Drug binding site. Magnified, orthogonal views of the additional central density in map (a) – blue mesh, and the fitted drug (iva), which is surrounded by residues in TM helices 6,10 & 12. The 2,4 *tert-*butyl groups at one end of the ivacaftor molecule allowed its orientation within the rocket-shaped density to be discriminated as described in Methods. F339 and Q343 within TM6 appear to form π-π and H-bonding interactions with the quinoline group at the other end of the drug. Small (glycine) residues at 868 and 985 in TM helices 10 and 12 may be important for accommodating the bulky *tert*-butyl groups whilst A981 may also satisfy hydrophobicity requirements for these groups. H-bonding with E871 by the phenolic hydroxyl group of ivacaftor is indicated. K185 in TM3 extends up into the central cavity to within 10Å of this group.

**Figure 3:**
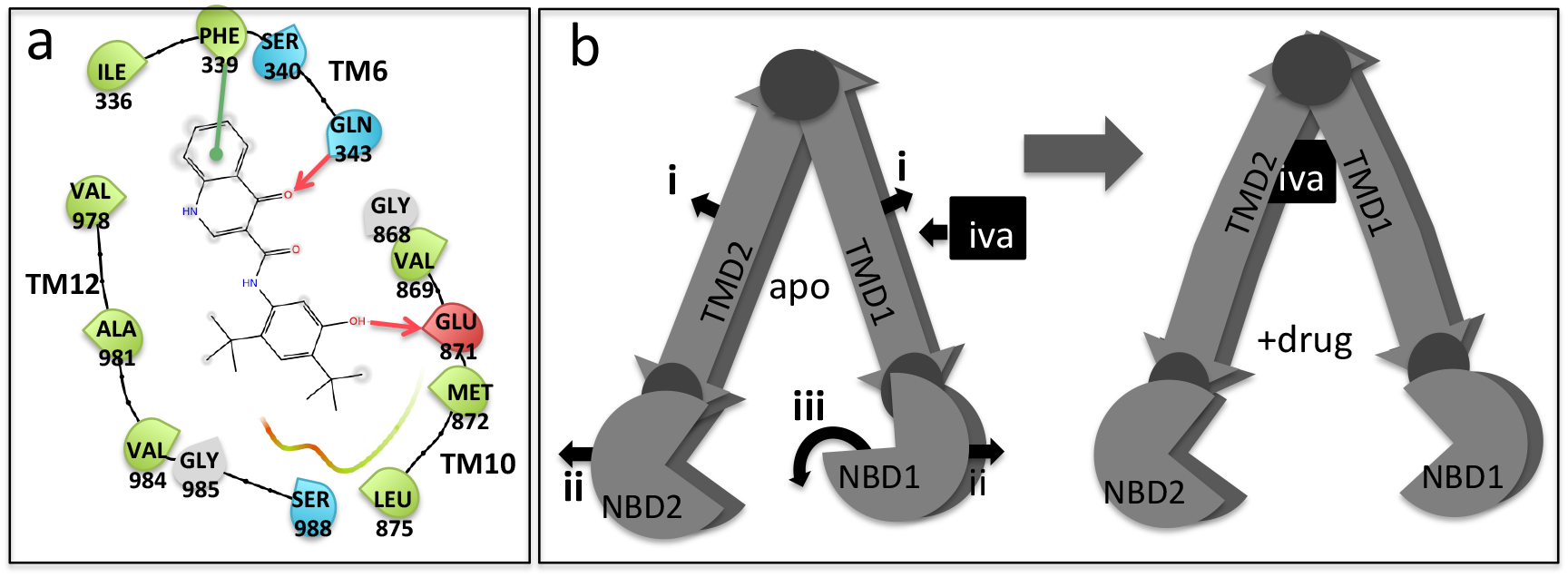
Ivacaftor binding and allostery. (a) Ligand interaction diagram of ivacaftor within the binding cavity of ABCB1. A threshold of 4.5 Å distance to neighbouring residues was used. H-bonding and π-π stacking are represented respectively with pink and green arrows. Negatively charged, hydrophobic and polar amino acids are coloured in red, light green and light blue, respectively. Residues F339 and Q343 appear to be close enough to form π-π and H-bonding with the quinolone group and E871 within TM10 with may H-bond with the phenolic hydroxyl of ivacaftor. (b) Cartoon representation of induced fit with ivacaftor binding to ABCB1. Left shows the apo state and the movements to the TMDs and NBDs needed for drug binding: (i) The TMD helices bow outwards and the central cavity enlarges. (ii) The NBDs become more separated. (iii) NBD1 rotates downwards. The small darker circles indicate hinge points between domains. Right shows the drug-bound state.

### Conformational changes associated with ivacaftor binding

The above results suggest that ivacaftor binding is associated with a wider separation of the two halves of ABCB1, similar to observations for the co-crystal structures of murine P-glycoprotein with the marine pollutant BDE-100(27–29). We examined whether any other local changes in individual domains could be identified as being specific to map *a* versus the other 4 maps. We assumed that maps *b* to *e* were representative of multiple conformations of the *apo* state). When we compared map *a* and its atomic model to the one most closely matching it: map *d (rmsd =1.17Å for 1018 atom pairs within 2Å, 1.46Å over all 1182 atom pairs)*, the local accommodation of the drug within the aqueous central cavity appeared to be associated with a local bulging outwards of the transmembrane helices (see Figure 3b). TM helices 11 and 12 seemed to move the most, perhaps because the residues in these helices are involved in interactions with the bulky *tert*-butyl groups of ivacaftor. When we compared map *a* with map *e* (which was the most different – *rmsd 1.40Å over 519 atom pairs within 2Å, 2.64Å over all 1182 atom pairs*) the outwards bulging of the TM helices upon ivacaftor accommodation was more noticeable. In both studies, the bulge of the TMDs results in a wider separation of the NBDs. NBD1 also showed a downward, shovel-like rotation away from the TMDs (Figure 3b). The net result of this was a downward (cytoplasmic) shift of the Walker A and B regions relative to intracellular loops 1 and 4 that connect NBD1 from the TMDs. This opens up the ATP binding site in NBD1. Similar changes, but with a smaller magnitude were also observed for NBD2. Whether the observed conformational shifts in the NBDs explain the increased ATPase activity of ABCB1 upon binding of a transport substrate remains to be determined by other means such as mutagenesis.

### Ivacaftor binding and comparisons with CFTR

Structural alignment of human CFTR with the atomic model generated for map *a* was carried out using the 5uak inward-facing structure of CFTR(30). The marginally narrower separation of the NBDs in the 5uak structure necessitated the separate rigid body alignment of the two arms of the CFTR structure with the equivalent regions of ABCB1 (see Methods). Prior to this, removal of the CFTR specific N-terminal ‘lasso’ structure was carried out. Other non-canonical aspects of the CFTR structure aligned poorly(19,30,31), in particular TM7 (31) and the extracellular side of TM8, which has a major deviation from helical structure before residue 936 (30)(Figure 4). Notably, CFTR TM10 was also significantly displaced versus the equivalent ABCB1 helix (Figure 4). The position of the ABCB1-fitted ivacaftor molecule relative to the ABCB1-aligned CFTR model is between TM helices 6 and 12 of CFTR (Figure 4, b&c). However the displacement of TM10 in CFTR results in that helix being much more distant from the ivacaftor position compared to in ABCB1. There is also no equivalent to K185 in TM3 of the aligned CFTR, however R352 would be close enough (6Å) to have an influence on the ivacaftor phenolic hydroxyl group. The ivacaftor molecule falls in a region of CFTR that has been associated with the ion channel gate and ion selectivity (32–38). It is clear from Figure 4 that the drug could not bind to CFTR in this position unless the clashing residues Q1144, R347 and R352 adopted different rotamers compared to those displayed in the *5uak* model. Whether CFTR in the inward-facing state can bind ivacaftor in the manner shown in Figure 4 is unknown, but if so, rotameric rearrangements of residues in TM helices 6 and 12 might be associated with channel opening and potentiation by ivacaftor. Well-characterised CF-causing mutations R347H, R352Q and R352W show low channel activity(39), and it is known that R347H and R352Q can be stimulated by ivacaftor (40). Conversely, R347P CFTR has undetectable levels of channel activity that could not be potentiated (39,40), suggesting that a rigid proline residue at this position may prevent channel opening, even in the presence of ivacaftor. In the outward-occluded conformational state of CFTR (PDBID 6o2p and 6msm), a strikingly asymmetric channel exists in one half of the transmembrane region formed by TM helices 1,2,3,6,11,12, see Figure 4d,e (19,41). The other half of the TMD (formed by TM helices 5,6,7,8,9,12 does not contain any obvious channel, largely because of the non-canonical configuration of TM helices 7, 8&9. In these occluded structures, residues R347, R352, Q1141 in TM6 and 12 that were discussed above do not form any obvious restriction to the channel along the direction of the membrane normal. This is difficult to rationalize in the light of the biochemical data and the effects of mutation of these residues. In the same state, ABCB1 shows a more symmetrical arrangement in the TMDs and a bi-lobed central cavity ((12), Fig. 4f). If the above mentioned residues do represent a channel gate, then one might propose that they represent a lateral gate, communicating between one lobe of the TMD and the other.

**Figure 4:**
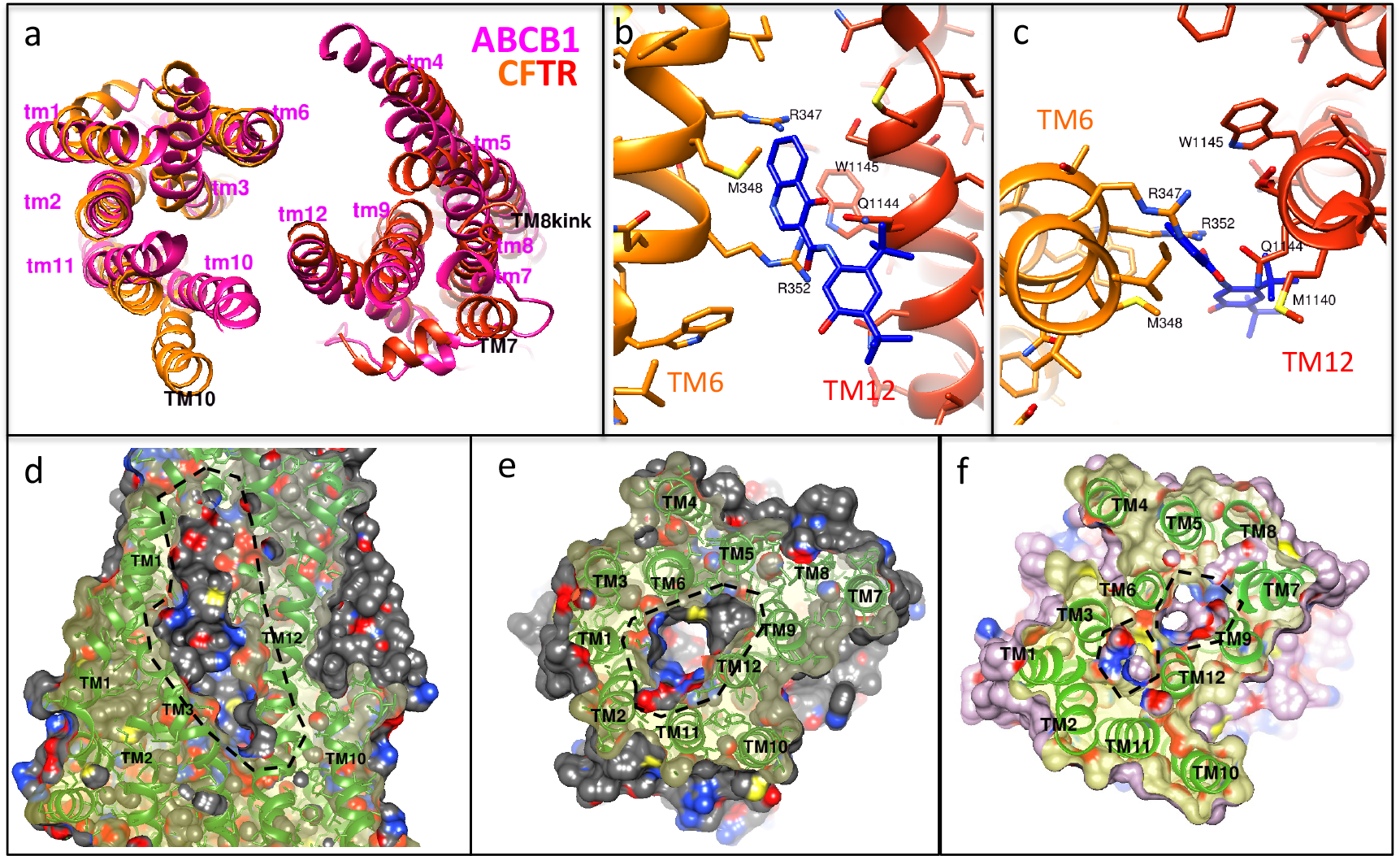
ABCB1-CFTR comparisons. (a) Alignment of the N- and C-terminal arms of CFTR in the inward-facing state (PDBID 5UAK, orange and red ribbons, respectively) with the equivalent regions of the ABCB1 structure represented by map *a* (pink ribbons). ABCB1 transmembrane helices are identified in lower case/pink text. CFTR helices differing significantly from the positions of the ABCB1 equivalents are indicated in upper case/black text. (b and c) When the ABCB1-fitted ivacaftor molecule (blue) is superimposed with the aligned CFTR model, it falls in a region associated with the channel gate and ion selectivity. Ivacaftor could not bind to CFTR in this position unless clashing residues Q1144, R347 and R352 adopted different rotamers compared to in the 5UAK model. Panel b shows the region viewed along the membrane plane, panel c from the extracellular side (similar to panel a). (d,e) Orthogonal slices through the transmembrane region of the outward/occluded state of CFTR (PDBID 6MSM), showing the asymmetric channel (dashed line). (f) Equivalent slice to panel e, but for the equivalent outward-occluded state of ABCB1 (PDBIB 6C0V) which displays a much more symmetrical profile and a bi-lobed central channel (dashed line).

## Methods

### Murine ABCB1a expression

*P. pastoris* harbouring the *opti-mdr3* gene were kindly provided by Prof. Ina L. Urbatsch, Texas Tech University (42). Cell culture was performed using the shake-flask method described by Beaudet and co-workers with minor modifications (43). Overexpression of ABCB1a was initiated by an addition of methanol and the expression was prolonged following an optimised procedure (13). Cell pellets were collected and stored at −80°C.

### Protein extraction and purification

Cell rupture was conducted using glass bead beating and microsomal membranes were prepared as described in (4). Microsomal membranes were diluted to 2.5 mg/ml prior to solubilisation in detergent-containing buffer (50 mM Tris pH 8.0, 10% (v/v) glycerol, 50 mM NaCl, 1 mM 2-mercaptoethanol and 2% (w/v) DDM). Solubilised materials were processed through multi-step protein purification described previously (13,14). Purified protein was concentrated to 5-10 mg/ml using a Vivaspin concentrator with 100-kDa cut-off, flash-frozen in liquid nitrogen and stored at −80°C.

### Drug binding assay

Drug binding and ABCB1a thermostability were assayed in three ways using an UNCLE (Unchained laboratories) instrument: Concentrated ABCB1a protein (1μg per sample) was diluted into 10μl of buffer (100mM Tris-HCl pH 8, 150mM NaCl, 10% glycerol, 0.1% DDM and 0.02% CHS) containing different concentrations of ivacaftor. The experiments were also repeated with the further addition of 100ng of CPM dye (7-Diethylamino-3-(4’-Maleimidylphenyl)-4-Methylcoumarin) which acts as a reporter of solvent-exposed cysteine residues (14,44,45). The mixtures were loaded into capillaries at 4°C and the latter were inserted into pre-chilled 16 well UNCLE capillary cassettes. Fluorescence emission spectra were recorded between 200nm and 700nm with an excitation wavelength of 266nm. Static light scattering was recorded simultaneously using the 266nm laser source. Data was gathered between 16°C and 90°C with a stepwise increase in the temperature (2°C increments) followed by the recording of spectra for all the samples. The total heating run required approximately 70min. Raw data was exported and then analysed for tryptophan fluorescence, static light scattering and CPM florescence using the GraphPad Prism software package.

### Specimen preparation for cryo-electron microscopy and data collection

Murine ABCB1a was initially buffer-exchanged into detergent/glycerol-free buffer. The protein was diluted to 1.1 mg/ml and incubated with 2 μM Ivacaftor (stock: 100 μM in DMSO) for 30 minutes on ice. Quantifoil 200 or 400 Au grids with a 1.3 micron spacing pattern were pre-treated prior to protein deposition as described previously (13). Briefly, grids were surface-cleansed by multiple chloroform washes to eliminate hydrophobic residues, then glow-discharged. A FEI Vitrobot MkIV was employed to facilitate sample vitrification where 3 μl of protein was gently loaded onto the middle of the grid, immediately blotted for 6 seconds and flash-frozen in liquid ethane. Grids were assessed for specimen quality (i.e. ice thickness, particle distribution, etc.) using a Polara G30 before shipping to the eBIC UK National facility for high resolution data acquisition on a FEI Titan Krios G2 microscope. Data were recorded at 300 kV using a 20 mV energy filter and a Gatan K2 electron detector. Images were acquired *via* the Volta phase plate (46) with phase shift range between 0.15 and 0.8 radian and fixed defocus at −0.5 μm. A total of ∼64 e^−^ Å^−2^ were used over 40 frames spanning 10 seconds of exposure and early and late frames were discarded. The magnification was calibrated at 1.043 Å/pixel. 2436 movie stacks were collected, and beam-induced image shift was eliminated using MotionCorr2 (47).

### Cryo-EM data processing, model building and refinement

The data processing of the 2436 images was entirely performed with *cis*TEM (48). A total of 1030 images were eliminated for possessing poor CTF fitting profiles and exceeding defocus values. The final set of 1406 images was divided into two similarly-populated subsets (696i and 710i) to independently assess the reproducibility of the 2D classification procedure. *Ab-initio* 3D reconstruction was performed employing ∼50,000 particles derived from several 2D classification runs of the 696i subsets. For each of the 5 classes that were generated *ab-initio*, a PDB model was derived using Molecular Dynamics Flexible Fitting (MDFF) with NAMD 2.12 as detailed previously (13). These models were used as a measure for comparing NBD separation in the five classes. *Ab-initio* 3D maps were then low pass filtered to 10Å and employed as starting templates for 3D refinement using the full final set of 104,000 selected particles. After full 3D refinement of the maps, real-space refinement of atomic models against the maps was performed within the PHENIX package (23) with the MDFF-derived models used as the initial inputs. Half-maps and Fourier shell correlation curves were generated employing *cis*TEM and the *calculate_fsc* routine. Local resolution was also estimated using the program ResMap (49). Rigid-docking of ivacaftor was performed with the fit-in-map routine of UCSF Chimera (50) and subsequently minimized in Schrodinger’s Maestro (Schrödinger Release 2021-2: Maestro, Schrödinger, LLC, New York, NY, 2021.) concomitantly allowing the rotamer adjustments of neighboring residues. The final result was verified to maintain the occupancy of the cryo-EM density.

### ABCB1/ABCC7 structural alignment

Residues 70-208,328-643 and 1014-1121 were selected in the atomic models of human CFTR and were aligned with the (mostly) N-terminal arm of ABCB1 using the ‘Matchmaker’ routine in Chimera(50). Residues 209-327,844-1013 and 1122-1480 were separately selected for alignment with the (mostly) C-terminal arm. As expected from the higher sequence homology, excellent alignment of the NBDs was observed, whilst most of the TM helices also aligned well.

## Availability of data and materials

Experimental density maps (*a*),(*e*) and the atomic models can be downloaded via the electron microscopy database (EMDB) and via the Protein Data Bank (RCSB) under the codes EMD-13059, 7OTG (map *a*), and EMD-13060, 7OTI (map *e*).

## Acknowledgements

We would like to thank Drs Daniel Clare and Alistair Siebert (eBIC) for support and advice for data collection and image processing. Access to the UK eBIC facility was via Block Allocation Grouping 16619. We thank Dr Hao Fan (A*STAR Bioinformatics Institute, Singapore) for advice and guidance. NT was supported by the Development and Promotion of Science and Technology Talent Project (DPST), the Institute for the Promotion of Teaching Science and Technology (IPST), Thailand. AB is supported by a University of Manchester/ASTAR Singapore PhD studentship. TS is supported by the Punjab education endowment fund (PEEF) with a Chief Minister Merit Scholarship (SSMS).

## Supplementary Figures

Supplementary Figure 1: ABCB1 affinity for ivacaftor measured by thermostability changes: (a) Ivacaftor binding detected using the ABCB1 intrinsic tryptophan fluorescence peak wavelength. A broad protein unfolding transition between 30-50°C was detected and ivacaftor binding exerted a detectable effect on the area under the curve (AUC) in this range. (b) Static light scattering (SLS), measured simultaneously to the tryptophan fluorescence showed a shift from increasing to decreasing light scattering, with transition around 30-50°C that was sensitive to ivacaftor. Arbitrary units are displayed for SLS. (c) CPM binding to ABCB1 cysteine residues detected an unfolding transition between 30-50°C that was sensitive to ivacaftor. Arbitrary units are displayed.

Supplementary Figure 2: (a) CTF correction and initial selection of images: Histogram of CTF fitting precision which determines the maximum resolution obtainable from an image. From the 2436 images 217 micrographs were removed to the right of the black bar (6 Ångstroms resolution or worse). A subset of images with thick ice gave very poor CTF estimation and were arbritrarily assigned a max resolution of 25Å. (b) Step-by-step workflow with sorting of images based on their CTF estimation and data processing of the remaining set and subsets.

Supplementary Figure 3: Cryo-EM images initially visualized with the CisTEM software suite. Left: Very high contrast particles can be readily observed due to the use of the Volta phase plate. The defocus used to record the image was ∼590nm. Particles were picked and later padded into a final box size of 200 pixels as shown on the right. The defocus of the exemplar image (right) was ∼780nm.

Supplementary Figure 4: Initial classification of the automatically-picked particles: (Left) A preliminary reference-free classification of 317,000 particle projections was carried out and 13 out of 60 classes that showed the expected features of ABCB1 were selected, corresponding to 108,000 particles. These were subsequently re-classified allowing up to 100 classes (right). Particles in a few remaining classes that had poorly defined features were removed, leaving a final dataset of 104,000 particles.

Supplementary Figure 5: 3D classification of particles. *Ab-initio* 3D classification of a subset representing about half of the ABCB1 dataset and allowing up to five 3D classes. The procedure superficially separated the particles into classes with wider or narrower separation of the NBDs. At this early stage of the procedure the 3D maps were relatively noisy (e.g. class c). Bottom right shows the number of particles used for each *ab-initio* class (blue); their percentages (green) and their distribution as a pie chart.

Supplementary Figure 6: Assessment of map *a* using a comparison of the two half-maps and the Resmap program. Upper panel shows the local resolution using colouring as indicated by the key on the right. Resolution is worse on surface-exposed regions and better for the TM helical region. The density for ivacaftor is indicated in the TM region slice (right, dashed line). The resolution of the map in this region is around 5.5Å. The histogram at the bottom shows the number of voxels within the map that fall in regions associated with a specific spatial resolution. The lowest resolution bar in each histogram (far right) shows all voxels in the map from regions with resolution equal to or worse than 9Å.

Supplementary Figure 7: Different assessments of the resolution of all five maps. (a) The histograms indicate the spread of resolution/variance across different regions of the maps using the Resmap program and comparing half-maps, each calculated wth half the particle numbers of the full maps. The number of voxels within 0.5Å resolution bands is plotted. The lowest resolution bar in each histogram (far right) shows all voxels from regions with resolution equal to or worse than the given resolution. (b) Resolution assessment using the Fourier shell correlation (FSC) calculated between the half-maps. (c) FSC plots for the relevant fitted model versus the full map generated by the Phenix package. For reference, the table lower right reproduces Table I in the main text and summarises all the data for resolution assessment of the maps.

Supplementary Figure 8: Comparison of model-to-map fits for different regions of map *a*. Upper panels show transmembrane (TM) helices 2,3 and 6 (green ribbon trace depicts secondary structure, ball and stick representation of side-chain atoms) with the map (mesh) coloured red within 3Å of any model atom, yellow if not. Viewing direction is along the membrane plane. Middle panels show the NBD domain interfaces as viewed from between the NBDs. Colouring as previously but the intracellular loops formed by the cytoplasmic portions of TM helices are indicated with the blue ribbon. The lower panels show the helical sub-domain of the NBDs. The right hand panel shows the correlation coefficient (CC) calculated between individual residues and the local map density. The dashed line indicates the mean CC value. The TMD residues display slightly higher CC than the NBDs. Loops between secondary structural elements show the worst correlations, especially extracellular loops (ECL) 1,2,4&5.

