## Supplemental Figures for "Structure of ABCB1/P-glycoprotein bound to the CFTR potentiator ivacaftor"

### Supplementary Figure 1

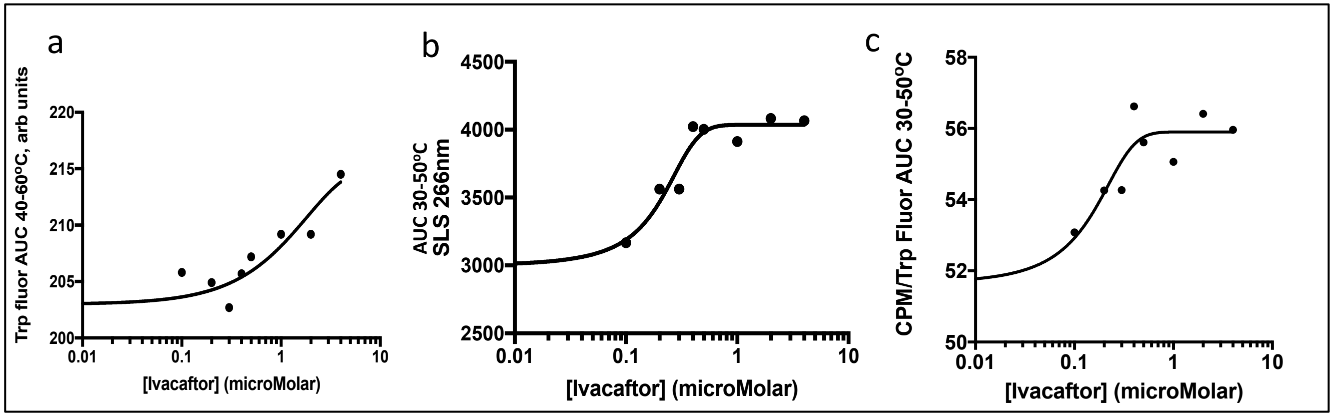

Supplementary Figure 2

### Initial processing of phase plate data

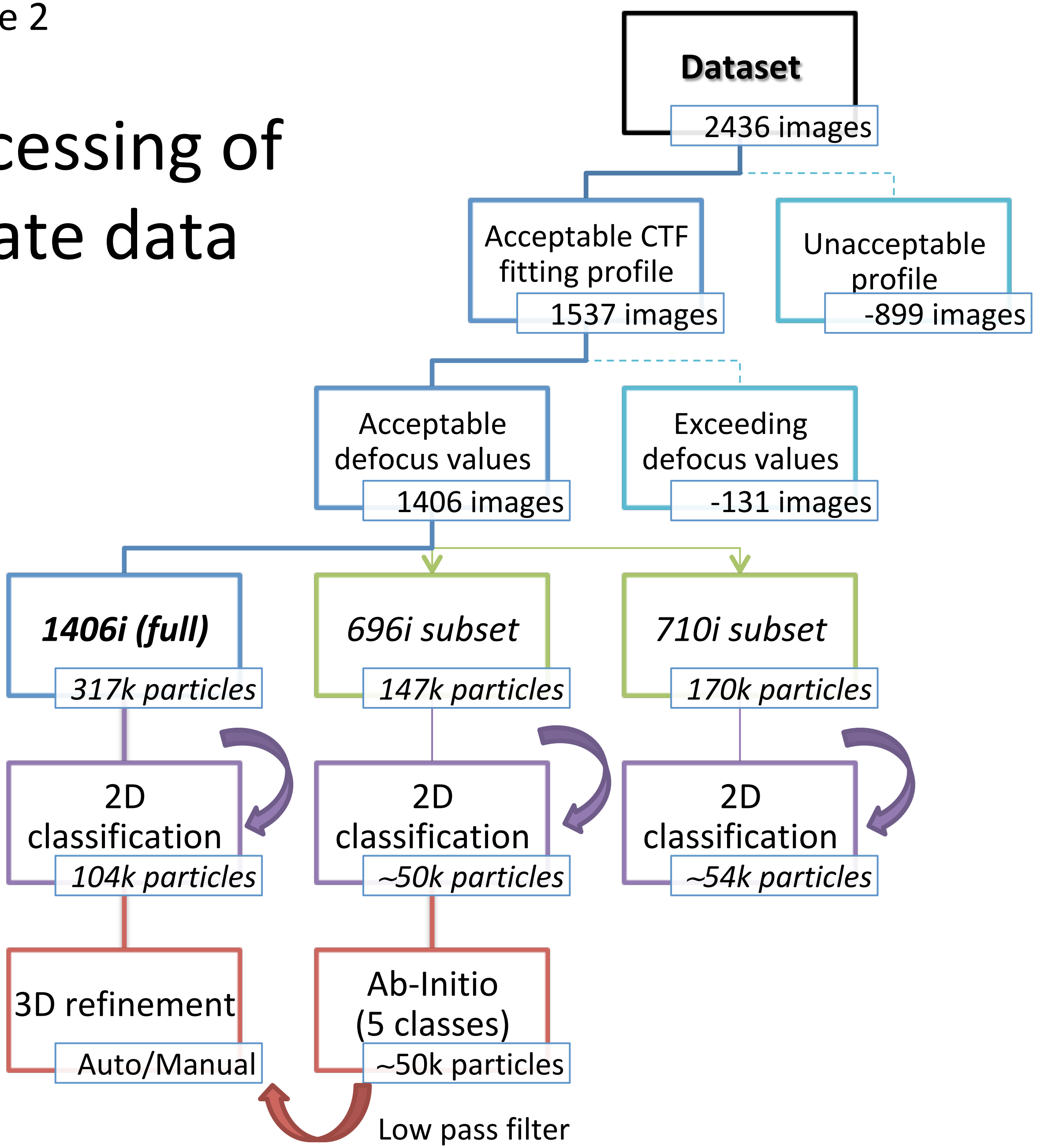

Supplementary Figure 3

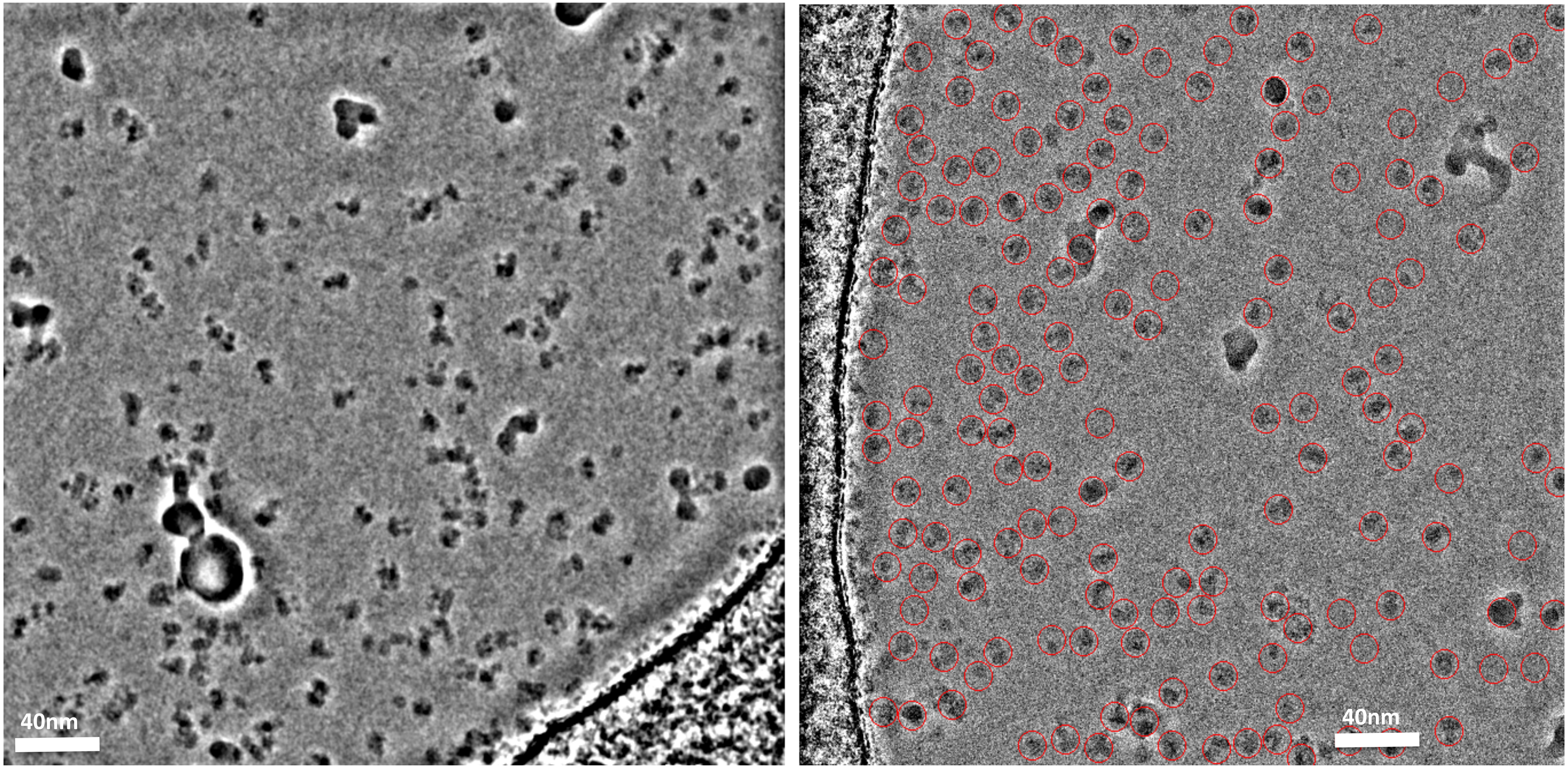

### Supplementary Figure 4

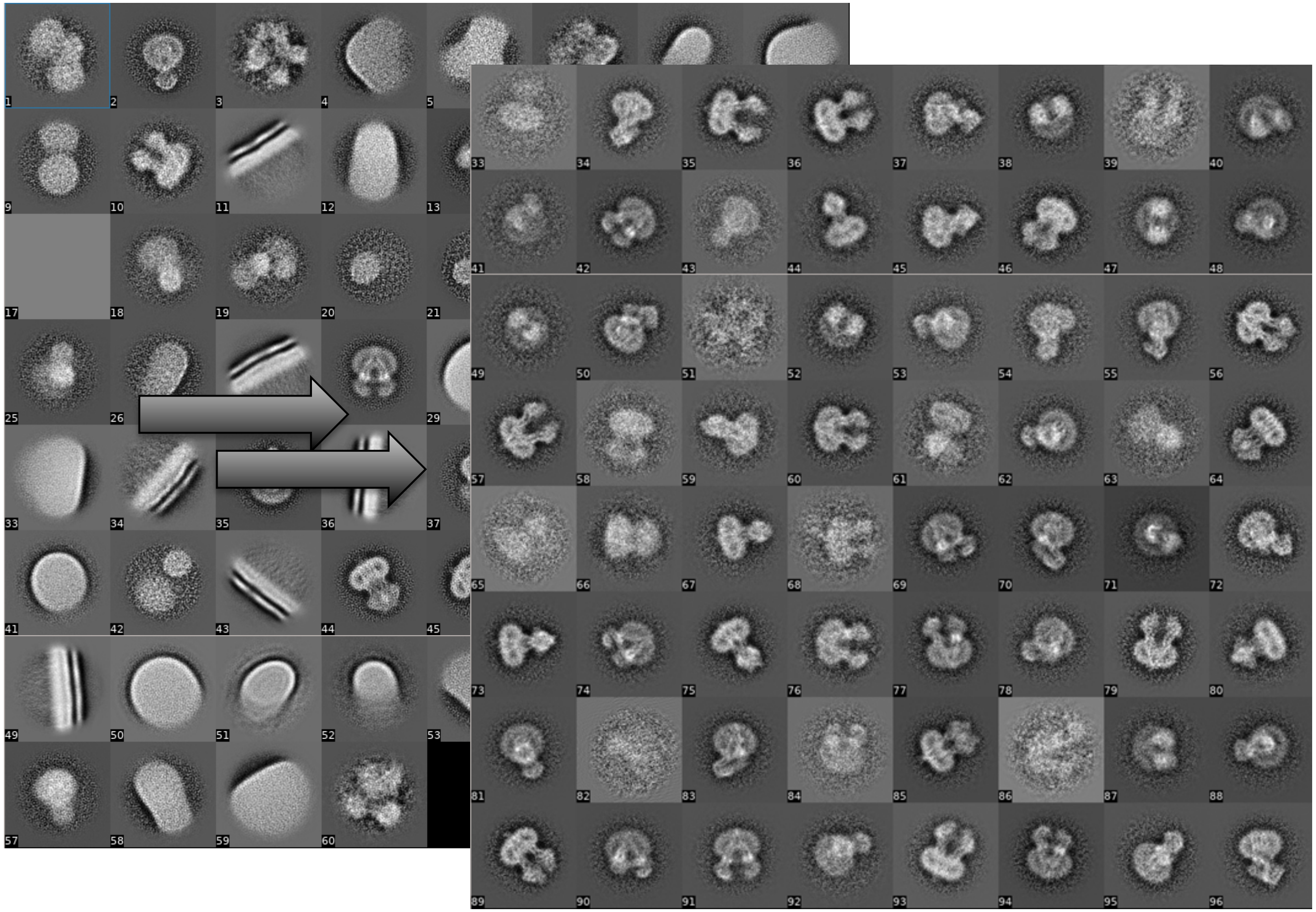

317k particles → 108k particles

Supplementary Figure 5

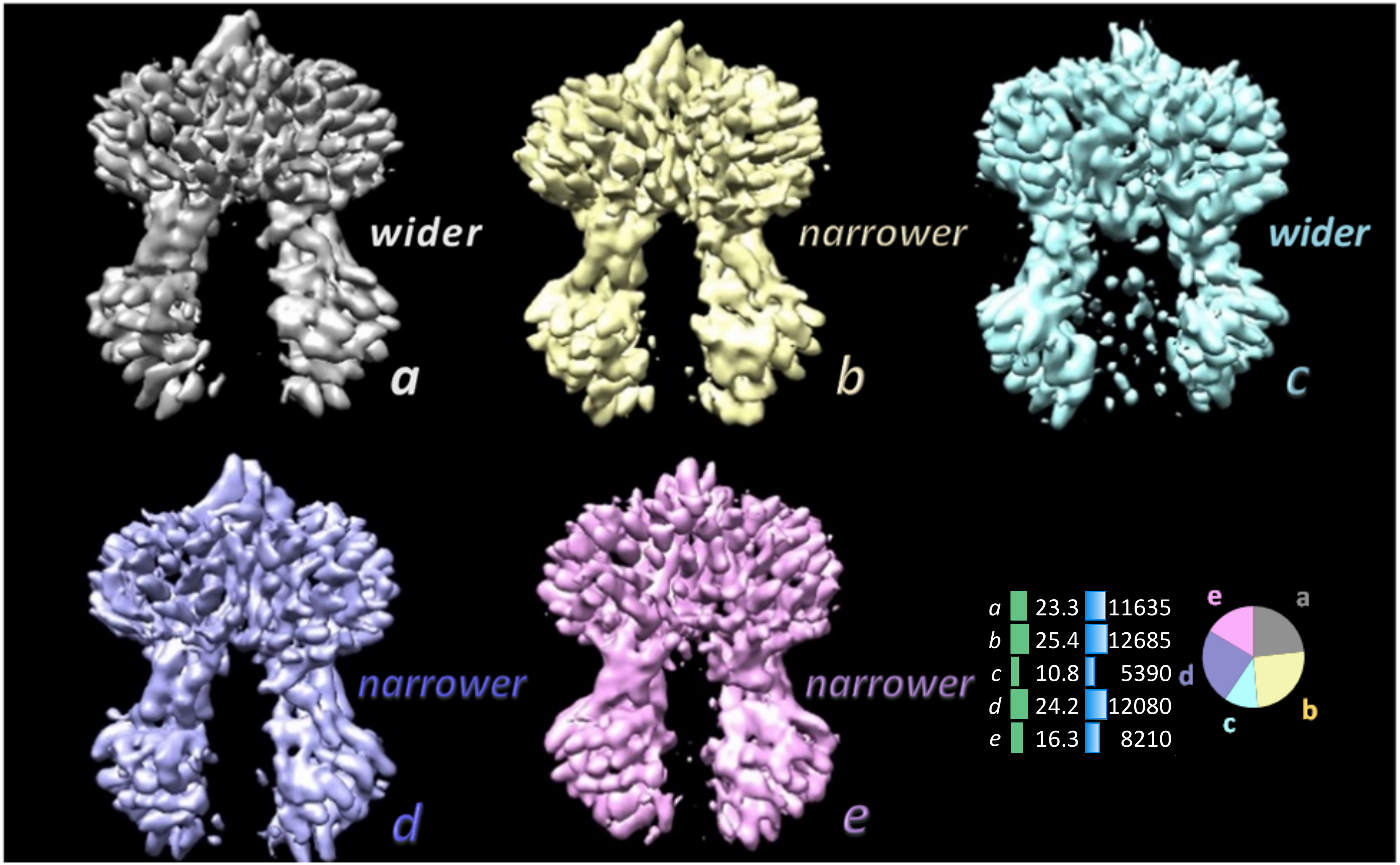

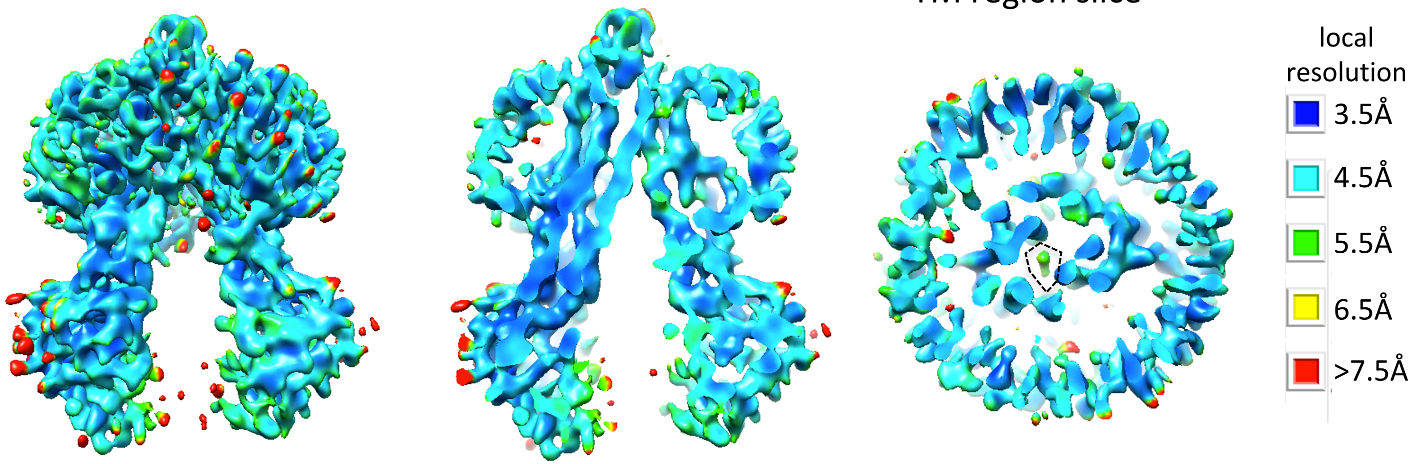


Colouring of different regions of map *a* according to local resolution (Resmap)

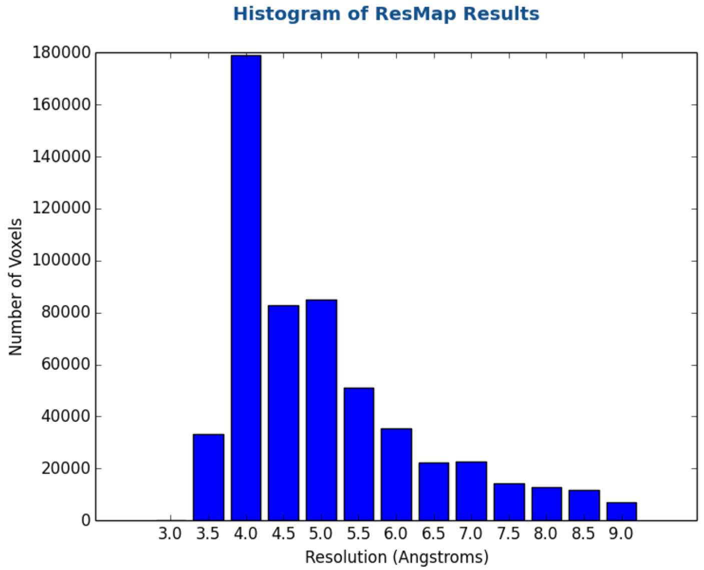


Resmap – Map *a*

Supplementary Figure 6

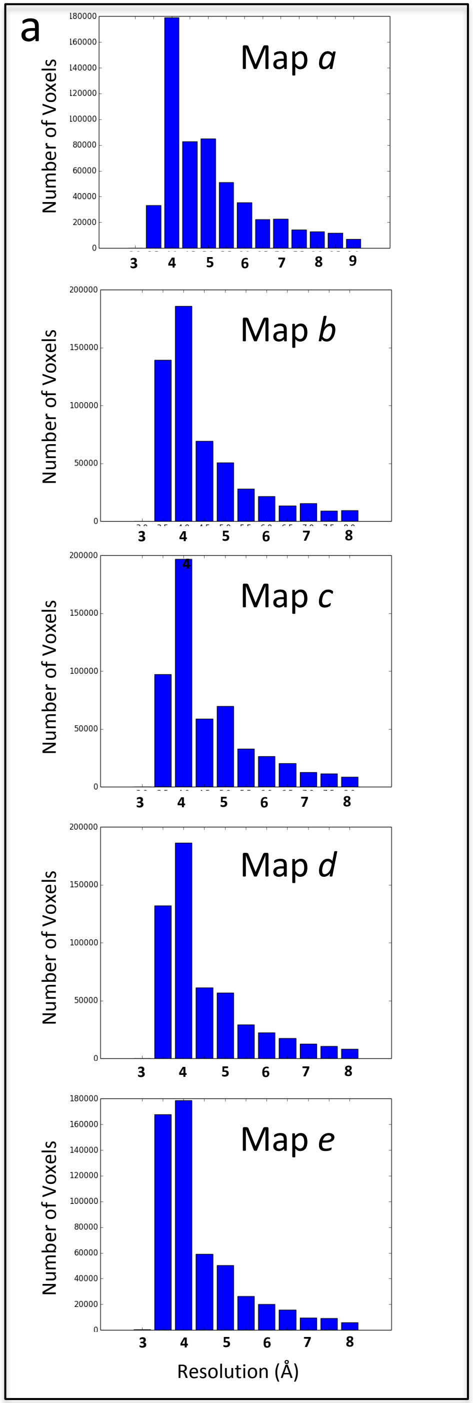


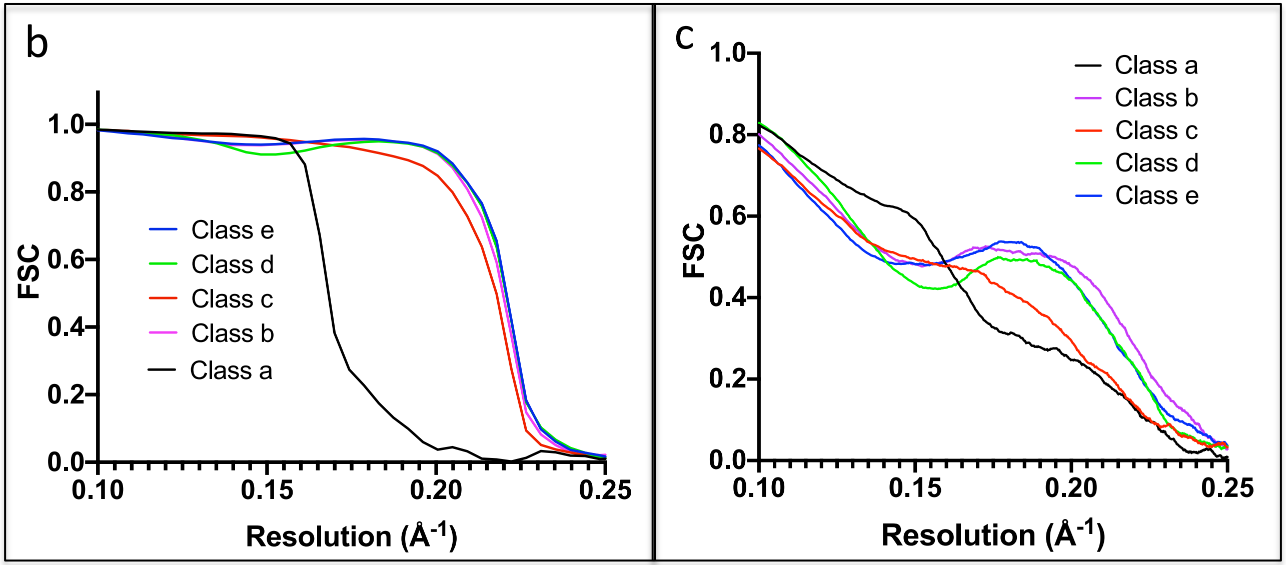


| Parameter | Map *a* | Map *b* | Map *c* | Map *d* | Map *e* |
|---|---|---|---|---|---|
| Global Resolution (Å, FSC=0.143) from unmasked half maps (CisTEM) | 5.4 | 4.3 | 4.3 | 4.2 | 4.2 |
| Global Resolution (Å, FSC=0.143) from map to model fit (Phenix) | 4.6 | 4.4 | 4.5 | 4.4 | 4.4 |
| Resolution (Å) from masked half maps (Mean,Mode from Resmap) | 5.0, 4.0 | 4.5, 4.0 | 4.6, 4.0 | 4.5, 4.0 | 4.4, 4.0 |
| CC map to model fit (mean, from 1182 residues, Phenix) | 0.73 | 0.67 | 0.63 | 0.69 | 0.65 |

Supplementary Figure 7

### Supplementary Figure 8

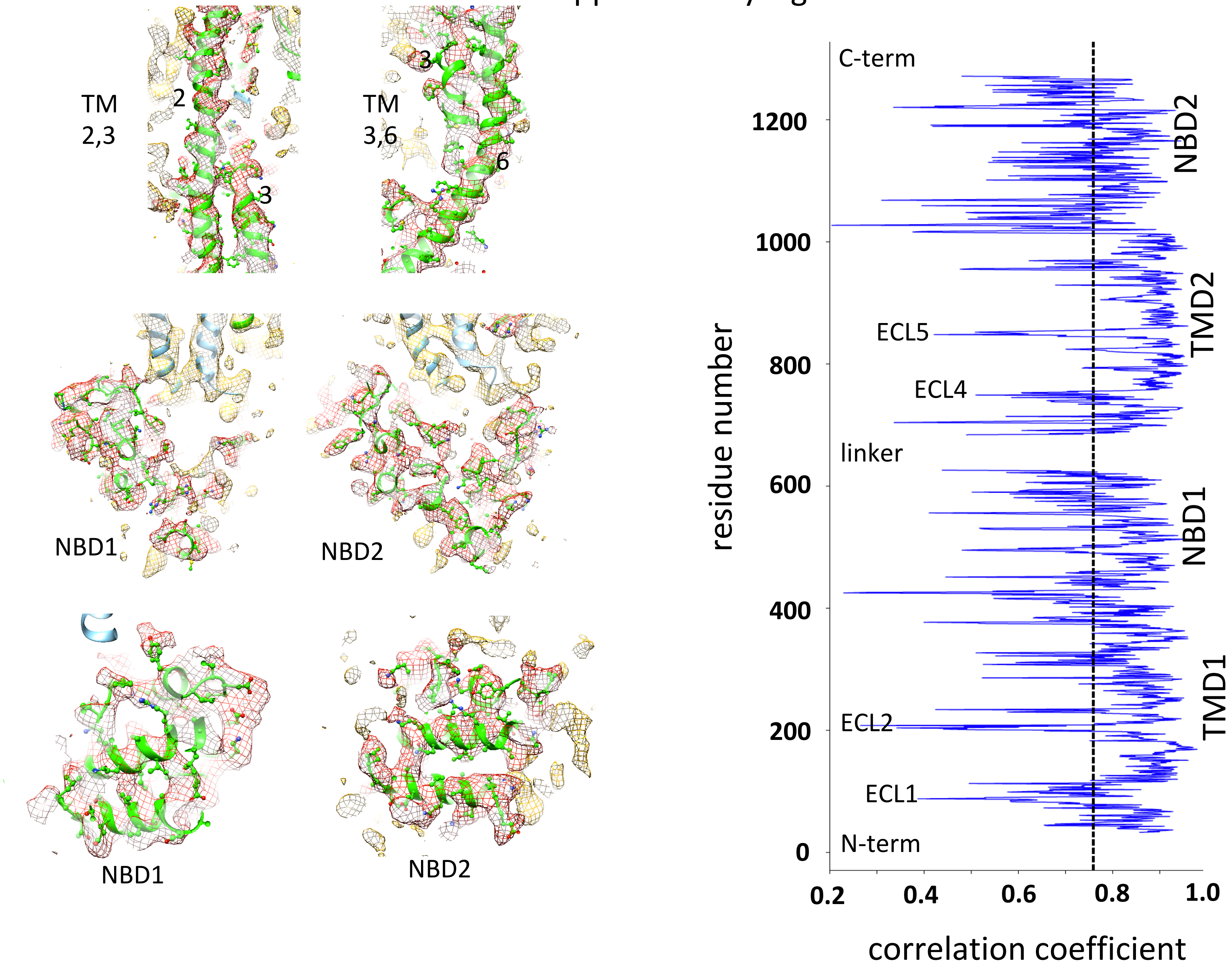
